# Distinct and shared contributions of diagnosis and symptom domains to cognitive performance in a case-control study of severe mental illness in the Paisa population

**DOI:** 10.1101/731265

**Authors:** Susan K. Service, Cristian Vargas Upegui, Mauricio Castano Ramírez, Allison M. Port, Tyler M. Moore, Marfred Munoz Umanes, Luis Guillermo Agudelo Arango, Ana M. Díaz-Zuluaga, Juanita Melo Espejo, María Cecilia López, Juan David Palacio, Sergio Sánchez Ruiz, Johanna Valencia, Terri Teshiba, Alesandra Espinoza, Loes Olde Loohuis, Juan De la Hoz Gomez, Benjamin Brodey, Chiara Sabatti, Javier I. Escobar, Victor I. Reus, Carlos Lopez Jaramillo, Ruben C. Gur, Carrie E. Bearden, Nelson B. Freimer

**Author notes:** co corresponding: Nelson B. Freimer, UCLA Center for Neurobehavioral Genetics, Semel Institute for Neuroscience and Human Behavior, Gonda Building, Room 3506, 695 Charles E. Young Drive South, Los Angeles, CA 90095, Carrie E Bearden, UCLA Center for Neurobehavioral Genetics, Semel Institute for Neuroscience and Human Behavior, A7-460 Semel Institute, 760 Westwood Plaza, Los Angeles, CA 90095.

## Abstract

**Background:** Severe mental illness (SMI) diagnoses display overlapping symptomatology and shared genetic risk, motivating trans-diagnostic investigations of disease-relevant quantitative measures. We analyzed relationships between neurocognitive performance, symptom domains, and diagnoses, in a large sample of SMI cases (ascertained agnostic to diagnosis) and healthy controls from a single, homogeneous population.

**Methods:** 2,406 participants (1,689 cases, 717 controls; mean age 39 years, 64% female) were assessed for speed and accuracy using the Penn Computerized Neurocognitive Battery (CNB). Cases carried structured-interview based diagnoses of schizophrenia (SCZ, n=160), bipolar-I (BP-I, n=519), bipolar-II (BP-II, n=204) and major depressive disorder (MDD, n=806). Linear mixed models, using CNB tests as repeated measures, modeled neurocognition as a function of diagnosis, sex, and all interactions. Follow-up analyses, in cases, included symptom factor scores obtained from exploratory factor analysis of symptom data, as main effects.

**Findings:** BP-I and SCZ displayed nearly identical impairments in accuracy and speed, across cognitive domains. BP-II and MDD performed similarly to controls, with subtle deficits in executive and social cognition. A three-factor model (psychosis, mania, and depression) best represented symptom data. Controlling for diagnosis, premorbid IQ, and disease severity, high lifetime psychosis scores were associated with reduced accuracy and speed across cognitive domains, while high depression scores were associated with increased social cognition accuracy.

**Interpretation:** Trans-diagnostic investigations demonstrated that neurocognitive function in SMI is characterized by two distinct profiles (BP-I/SCZ and BP-II/MDD), and is associated with specific symptom domains. These results suggest the utility of this design for elucidating SMI causes and trajectories.

## Introduction

Schizophrenia (SCZ), bipolar disorder (BPD), and major depressive disorder (MDD), the diagnoses that together constitute severe mental illness (SMI), are each among the largest contributors to the global burden of disease (1). The splitting of SMI into these diagnostic categories, based on symptoms and classical disease trajectories, has long dominated psychiatric research and clinical practice. Research from across the behavioral sciences has increasingly challenged these dichotomies, and stimulated efforts to reorient psychiatric research around systems of dimensional phenotypes (2).

Two main classes of dimensional phenotype have been proposed: (a) symptoms, (e.g. “psychosis”), which are components of specific diagnostic categories, but may be present across multiple categories, and (b) quantitative measures that assess neurobehavioral domains, such as cognitive function, that are outside of the current diagnostic framework and yet characterize SMI. Cognitive function is clearly impaired in SMI cases overall, relative to controls, and recent work has demonstrated that symptom components, such as depression and psychosis, and diagnosis have potentially additive effects on cognition (3, 4). Few large psychiatric case samples have obtained the measures needed to test hypotheses relating symptoms and cognition to SMI, and analyses have been limited by the heterogeneity across study samples in the approaches used for both ascertainment of participants and their phenotypic assessment. Furthermore, going forward, it remains unclear how use of dimensional phenotypes would impact understanding of the biological underpinnings of SMI, for example, through genetic studies.

Genome-wide association studies (GWAS) of SMI have already identified hundreds of loci that are unequivocally associated with these disorders (5). Most of these significant associations are to a specific diagnosis. However, analyses of the totality of genetic variation represented in these GWAS datasets indicate that the overall polygenic contribution to disease risk is largely transdiagnostic (6). Taken together, existing data indicate the need for study designs that both include and transcend categorical diagnoses, incorporating dimensional phenotypes that may be distinct to SMI subtypes and those that are shared across them.

We report here, in a large and uniformly assessed case/control cohort, our test of the hypothesis that neurocognitive performance is associated with both SMI diagnoses and trans-diagnostic symptoms. Four aspects of this study are, to our knowledge, unique. First, the availability of electronic medical records (EMR) from two psychiatric hospital systems in the Paisa region of Colombia enabled us to ascertain SMI cases agnostic to specific diagnoses and to incorporate measures of lifetime disease severity in our analyses. Second, the cohort includes large numbers of cases from each of the major SMI diagnostic categories, from MDD to SCZ. Third, we assessed in all cases a set of symptoms that would typically be probed, using structured interview branching logic, only in individuals that have responded positively to specific screening questions. In this manner we were able to assess, for example, symptoms associated with depression and mania that would not typically be queried of cases with SCZ. Finally, the study sample derives from a single population that is relatively homogenous genetically and culturally, thereby minimizing confounds due to inter-population variability.

## Methods

### Sample Ascertainment and procedures

Cases with SMI were ascertained through EMR at Clínica San Juan de Dios de Manizales (CSJDM) in Manizales, Caldas and the Hospital Universitario San Vicente Fundación (HUSVF) in Medellín, Antioquía, beginning in 2017 (Supplementary Figure 1). Individuals were invited to participate in the project based on the following criteria: (a) EMR diagnosis of a mood or psychosis spectrum disorder with a history of at least one inpatient hospitalization or treatment for symptoms considered sufficiently severe by a referring psychiatrist to warrant such hospitalization; (b) presenting symptoms were not clearly caused by a substance use disorder, in the judgment of an evaluating clinician; (c) have two Paisa surnames; (d) aged 18 or above; (e) understand and sign an informed consent document; (f) no intellectual disability, and (g) no history of serious brain trauma or neurological disorder. Analyses reported here include individuals diagnosed with SCZ, BP-I, BP-II, or MDD on structured interviews (see Study Measures).

Healthy controls were ascertained from the same communities as cases, and recruited from friends, neighbors, or in-laws of cases, or from university students/staff and hospital staff. All controls met the following criteria: (a) no (current or lifetime) SMI, as evaluated through the overview screening module of the NetSCID (b) no current substance use disorder, and (c) fulfillment of criteria c-g described for cases. Cases and controls were reimbursed for transportation costs but were not otherwise compensated.

Cases and controls were evaluated at the hospitals. Before performing any assessment and after verifying inclusion and exclusion criteria, all participants signed an informed consent form. All procedures were approved by the IRB of the University of Antioquia (Comité de Ética del Instituto de Investigaciones Médicas de la Universidad de Antioquía), the Hospital San Vicente Fundación, the CSJDM, UCLA, UCSF, and the University of Pennsylvania.

### Study Measures

#### Cases

Data collected included prior psychiatric contacts and hospitalizations, and medication history. We obtained lifetime DSM-5 diagnoses and cross-diagnosis symptom-level data through structured interviews using a Spanish translation of NetSCID, a computerized version of the Structured Interview for DSM-5 (7, 8). Use of this instrument, with built-in algorithms and decision trees for determining diagnosis, increases reliability and reduces branching errors that could lead to misdiagnosis. NetSCID modules for case assessment included: overview, screener of major psychopathology, mood disorders, psychotic disorders, and trauma and stressor-related disorders. To assess trans-diagnostic symptomatology, we administered to cases seven supplementary questions about specific symptoms of fatigue, grandiosity, decreased need for sleep, flight of ideas, hypersomnia, apathy and anhedonia.

#### Controls

To screen for psychopathology in potential control participants, we used the NetSCID overview module.

#### All participants

Data collected included demographic information, medication use, substance use, a brief assessment of current severity of a range of psychiatric symptomatology (the 45-question Symptom Assessment Questionnaire (SA-45) (9)), and the Word Accentuation Test (WAT), a reading test for Spanish speakers, based on irregular accentuation of words (10), that has been validated in that group as a measure of premorbid IQ (see Supplementary Table 1 for the sample size available for each measure).

To assess speed and accuracy of neurocognitive performance across five domains related to specific brain systems hypothesized to be most strongly associated with SMI (executive function, memory, complex cognition, social cognition, and motor speed), we used nine tests from the Penn Computerized Neurocognitive Battery (CNB) (11), a psychometrically well-validated online battery (Supplementary Table 2) with standardized automated QA and scoring procedures. The CNB has been validated across a wide age range, in both community samples and psychiatric populations (12, 13). Data for speed were multiplied by −1 so that poorer performance (longer response time) would result in a lower value. All evaluators were extensively trained by the Penn team using both web-based training modules, on-site training (by RCG and AMP), and web-based supervision. Age and education strongly affect neurocognitive performance; therefore, as is standard for CNB analyses (14), the raw data for each test were regressed on age, age^2^, age^3^, education, and an age by education interaction, and residuals used for further analyses (see Regression Models, below). Residuals were winsorized at the top and bottom 1% level to reduce the influence of extreme outliers and transformed to z-scores based on the mean and SD in control participants.

All study data were collected and managed using REDCap electronic data capture tools hosted at UCLA (15, 16).

### Regression Models of CNB Accuracy and Speed

Z-scores were modeled as a function of diagnosis, sex, test domain and all interactions using linear mixed models (LMM), with individual CNB tests as repeated measures, as in (14). Separate analyses were conducted for accuracy and speed, which show different factorial structures (17), to reduce the dimensionality of the analyses, as we had no hypotheses involving accuracy by speed interactions. Analyses were done using the R function lme() in the nmle package (18, 19).

### Assessment of Possible Confounding Effects of Medication Use

Self-report data were collected on all participants for usage of 30 psychiatric medications and grouped into three categories (Supplementary Table 3): antidepressants (15 medications), antipsychotics (11 medications) and mood stabilizers (four medications). In each category, a binary indicator of medication use was constructed for each participant. The LMM analysis of z-scores was repeated including these three covariates to assess robustness of conclusions to medication use effects.

### Factor Analysis of Symptom Data in Cases

Symptom data for cases were obtained from the NetSCID interview, and supplementary questions. Symptoms were considered present in cases if they were endorsed at any point in the NetSCID (i.e., lifetime) or in additional queries, and considered absent if they were never endorsed, and confirmed as absent at least once. A total of 40 symptoms were evaluated, and we retained for analysis symptoms endorsed by at least 2·5% of cases. Missing symptom data for cases from the NetSCID were imputed a single time using bootstrapped expectation-maximization (EMB) by the amelia() function in the R Amelia package (20).

We conducted an exploratory item-factor analysis (21) on the matrix of tetrachoric inter-item correlations using weighted least-squares extraction and promax rotation. The number of factors to retain was determined by a combination of the minimum average partial (MAP) method (22), parallel analysis (23) with Glorfeld correction (24), visual examination of the scree plot, and theory. MAP was implemented by the nfactors() function in the psych package (25) in R; corrected parallel analysis was implemented by the fa.parallel() function (also in psych). Visual examination of the scree plot involved subjective judgment of the point on the plot where the eigenvalues began to form an approximate linear trend. In an analysis that used only cases, we repeated the LMM described above, including symptom factor scores as covariates.

### Assessment of potential confounding effects of premorbid IQ and disease severity in cases

We evaluated the robustness of our conclusions regarding the effect of symptom factors on cognition by performing additional LMM analyses in cases. In one analysis, we added the WAT score in the model as a covariate, to control for effects of premorbid IQ on cognition. Prior to this analysis, raw CNB data were adjusted only for age, age^2^, age^3^, and not education.

In a second set of analyses, we controlled for effects of lifetime and current illness severity on cognition. As a proxy for lifetime disease severity, we extracted all available EMR data on the number of visits of each participant to the emergency department or inpatient unit of the CSJDM in Manizales and included this variable as a covariate (complete EMR data were not available from cases recruited at HUSVF in Medellín). As a measure of current illness severity, we used the Global Severity Index of the SA-45.

### Role of the Funding Source

The funders (The US National Institute of Mental Health) had no role in the study design; the collection, analysis, and interpretation of data, the writing of the report or the decision to submit the report for publication.

## Results

A total of 3,467 participants completed clinical assessments and were recruited into the study (Supplementary Figure 1). CNB data for 901 participants (817 cases and 84 controls) were missing (Supplementary Table 4). Among the remaining 2,566 participants (1,849 cases and 717 controls), 160 cases did not qualify for a NetSCID primary lifetime diagnosis of SCZ, BP-I, BP-II, or MDD, and were excluded from analysis. A summary of basic demographic information for each diagnostic category for the remaining 2,406 participants is in Table 1; controls did not differ from cases in terms of sex or years of education, but controls were significantly younger than cases.

**Table 1.** Demographic and medication information for 2,406 participants.

|  | Schizophrenia |  | Bipolar Disorder I |  | Bipolar Disorder II |  | Major Depressive Disorder |  | Controls |  | P-value comparing Cases to Controls |
| --- | --- | --- | --- | --- | --- | --- | --- | --- | --- | --- | --- |
|  | Male | Female | Male | Female | Male | Female | Male | Female | Male | Female |  |
| N | 132 | 28 | 207 | 312 | 56 | 148 | 210 | 596 | 268 | 449 | 0.467 |
|  | Mean | SD | Mean | SD | Mean | SD | Mean | SD | Mean | SD |  |
| Age | 37.4 | 12.8 | 44.1 | 13.8 | 38.7 | 14.4 | 39.3 | 14.1 | 36.7 | 14.4 | 2.71E-09 |
| Years of Education | 10.4 | 3.3 | 11.1 | 4.2 | 12.3 | 3.7 | 11.9 | 3.7 | 11.7 | 3.5 | 0.368 |
| Premorbid IQ (WAT score) | 27.1 | 8.6 | 29.9 | 10.0 | 31.9 | 8.3 | 31.1 | 8.1 | 29.9 | 8.3 | 0.208 |
| SA45 Global Severity Index | 38.6 | 34.6 | 30.7 | 31.0 | 55.2 | 39.8 | 53.0 | 37.9 | 12.5 | 13.8 |  |
| Number of hospitalizations and ER visits | 14.7 | 16.3 | 12.2 | 12.6 | 5.9 | 9.0 | 3.7 | 6.0 | NA | NA |  |
|  | Percent of Total | N | Percent of Total | N | Percent of Total | N | Percent of Total | N | Percent of Total | N |  |
| Percent on Antipsychotics | 89% | 142 | 68% | 353 | 52% | 106 | 33% | 264 | 0% | 0 |  |
| Percent on Antidepressants | 27% | 43 | 23% | 119 | 47% | 95 | 72% | 577 | 0% | 1 |  |
| Percent on Mood Stabilizers | 23% | 37 | 70% | 363 | 68% | 138 | 19% | 151 | 1% | 5 |  |
|  |  |  |  |  |  |  |  |  | NA | NA |  |
| Percent in bottom half of psychosis factor score | 0% | 0 | 20% | 102 | 54% | 110 | 80% | 646 | NA | NA |  |
| Percent in top half of psychosis factor score | 100% | 160 | 80% | 417 | 46% | 94 | 20% | 160 | NA | NA |  |
| Percent in bottom half of mania factor score | 64% | 103 | 2% | 11 | 8% | 17 | 89% | 715 | NA | NA |  |
| Percent in top half of mania factor score | 36% | 57 | 98% | 508 | 92% | 187 | 11% | 91 | NA | NA |  |
|  |  |  |  |  |  |  |  |  | NA | NA |  |
| Percent in bottom half of depression factor score | 93% | 149 | 68% | 354 | 33% | 67 | 46% | 367 | NA | NA |  |
| Percent in top half of depression factor score | 7% | 11 | 32% | 165 | 67% | 137 | 54% | 439 | NA | NA |  |

### Associations between Diagnosis and Cognitive Performance

Both accuracy and speed showed significant interactions of diagnosis and cognitive test domain, indicating that the diagnostic groups differed in their profile of cognitive deficits (Supplementary Figure 2, Supplementary Table 5A). For both accuracy and speed the four patient groups bifurcated into two profiles (Figure 1, Table 2), with SCZ and BP-I showing greater deficits than BP-II and MDD. The pattern of deficits was nearly identical for SCZ and BP-I, with greater deficits across executive function (where effect sizes neared and exceeded 1 SD), social cognition and motor speed tests relative to memory and complex cognition. While participants with SCZ tended to have less education than those diagnosed with BP-I (Table 1), the similarity of SCZ and BP-I profiles persists when cognitive data were not adjusted for education (data not shown). BP-II and MDD groups likewise had similar profiles, with more subtle deficits in executive functions (effect sizes 0.5 SDs), social cognition and motor speed, while performance in other domains was at normative levels.

**Figure 1.**
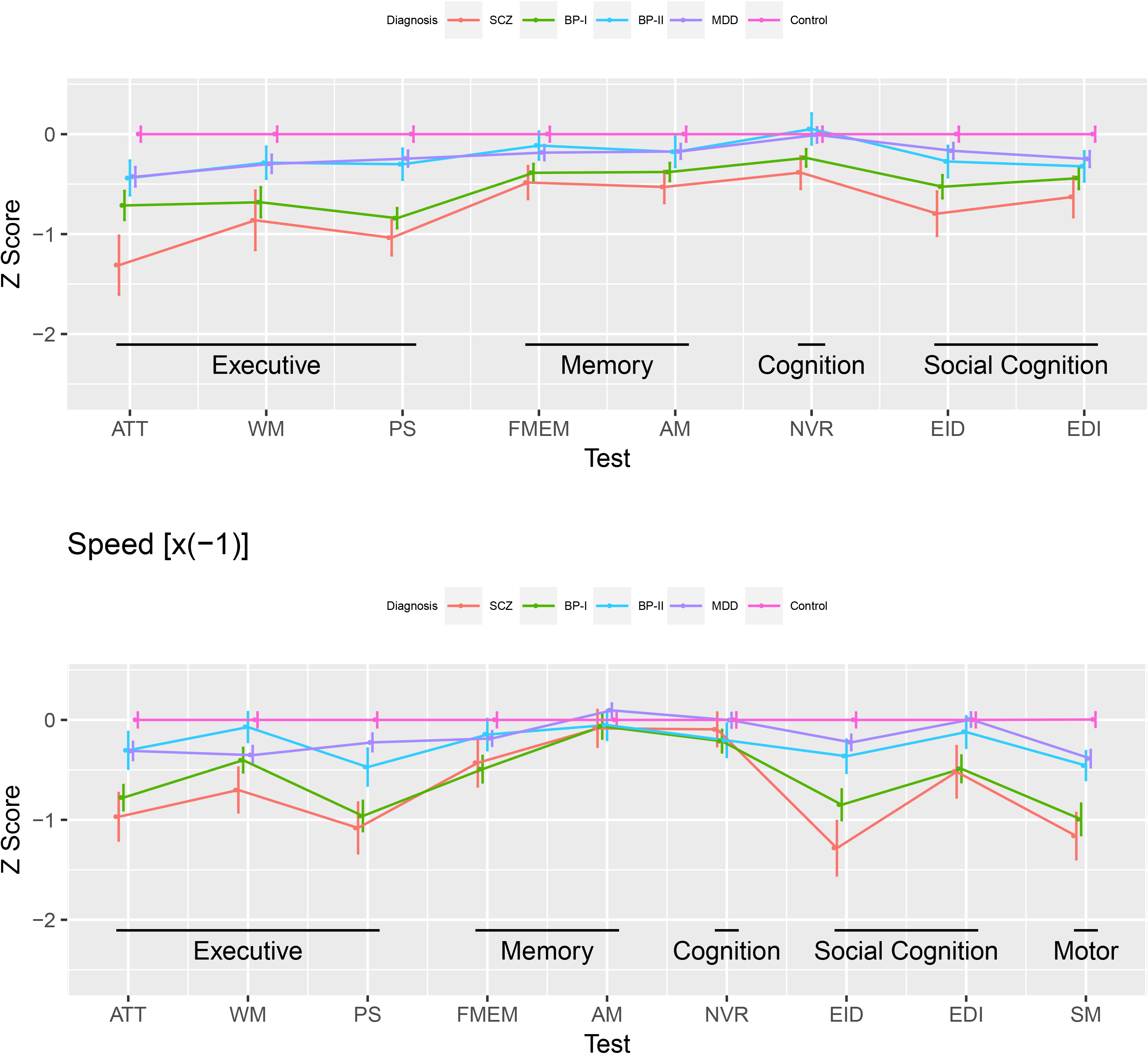
Z-scores on Accuracy (top) and Speed (bottom) profiles for tests assessing Executive Function, Memory, (Complex) Cognition, Social Cognition, and Motor Speed. Data for Speed were multiplied by −1 so that poorer performance (slower speed), would result in a lower value. Z-scores were generated relative to Controls (n=717). SCZ: schizophrenia (n=160) BP-I: bipolar disorder I (n=519), BP-II: bipolar disorder II (n=204), MDD: major depressive disorder (n=806). Test abbreviations: ATT = Continuous Performance Test; WM = Letter-N-Back test; PS = Digit Symbol Test, matching trials; FMEM = Face Memory test; AM = Digit Symbol test, recall trials; NVR = Matrix Analysis test; EID = Emotion Recognition test; EDI = Measured Emotion Differentiation test; SM = Motor Praxis test. Error bars are the 95% confidence intervals.

**Table 2.** Analyses of accuracy (A) and speed (B) z-scores for each Penn-CNB test. ATT = Continuous Performance Test; WM = Letter-N-Back test; PS = Digit Symbol Test, matching trials; FMEM = Face Memory test; AM = Digit Symbol test, recall trials; NVR = Matrix Analysis test; EID = Emotion Identification test; EDI = Measured Emotion Differentiation test; SM = Motor Praxis test. The reference category is control-females

| A<br>Coefficient |  | Executive Function |  |  | Episodic Memory |  | Complex Cognition | Social Cognition |  |
| --- | --- | --- | --- | --- | --- | --- | --- | --- | --- |
|  |  | ATT | WM | PS | FMEM | AM | NVR | EID | EDI |
| Intercept | Estimate | -0.019 | -0.067 | -0.023 | 0.004 | -0.025 | -0.093 | 0.081 | 0.023 |
|  | P | 7.29E-01 | 2.09E-01 | 6.04E-01 | 9.21E-01 | 5.44E-01 | 2.52E-02 | 9.06E-02 | 6.16E-01 |
| MDD | Estimate | -0.415 | -0.287 | -0.235 | -0.179 | -0.168 | 0.002 | -0.163 | -0.244 |
|  | P | 4.36E-09 | 2.52E-05 | 3.27E-05 | 8.64E-04 | 1.70E-03 | 9.77E-01 | 7.32E-03 | 2.77E-05 |
| BP-II | Estimate | -0.428 | -0.276 | -0.292 | -0.109 | -0.172 | 0.058 | -0.270 | -0.318 |
|  | P | 8.98E-05 | 8.73E-03 | 9.06E-04 | 1.90E-01 | 3.77E-02 | 4.84E-01 | 4.10E-03 | 4.35E-04 |
| BP-I | Estimate | -0.721 | -0.681 | -0.848 | -0.390 | -0.380 | -0.236 | -0.544 | -0.445 |
|  | P | 1.69E-19 | 7.19E-18 | 2.28E-38 | 1.58E-10 | 4.60E-10 | 1.10E-04 | 2.70E-15 | 1.54E-11 |
| SCZ | Estimate | -1.372 | -0.896 | -1.092 | -0.509 | -0.554 | -0.417 | -0.858 | -0.655 |
|  | P | 6.62E-27 | 7.16E-13 | 5.50E-27 | 8.20E-08 | 5.57E-09 | 1.15E-05 | 1.09E-15 | 1.91E-10 |
| Male | Estimate | 0.051 | 0.180 | 0.061 | -0.011 | 0.068 | 0.249 | -0.218 | -0.062 |
|  | P | 4.03E-01 | 2.44E-03 | 2.09E-01 | 8.06E-01 | 1.42E-01 | 8.15E-08 | 2.91E-05 | 2.15E-01 |

| B<br>Coefficient |  | Executive Function |  |  | Episodic Memory |  | Complex Cognition | Social Cognition |  | Motor |
| --- | --- | --- | --- | --- | --- | --- | --- | --- | --- | --- |
|  |  | ATT | WM | PS | FMEM | AM | NVR | EID | EDI | SM |
| Intercept | Estimate | -0.052 | -0.052 | -0.042 | -0.017 | -0.024 | 0.104 | 0.054 | 0.031 | -0.077 |
|  | P | 3.76E-01 | 3.65E-01 | 5.00E-01 | 7.58E-01 | 6.45E-01 | 3.99E-02 | 3.77E-01 | 5.77E-01 | 2.18E-01 |
| MDD | Estimate | -0.357 | -0.431 | -0.170 | -0.157 | 0.084 | -0.112 | -0.182 | 0.007 | -0.347 |
|  | P | 5.97E-06 | 1.66E-08 | 4.13E-02 | 3.51E-02 | 2.24E-01 | 1.03E-01 | 2.57E-02 | 9.30E-01 | 2.83E-05 |
| BP-II | Estimate | -0.398 | -0.082 | -0.442 | -0.181 | -0.072 | -0.310 | -0.363 | -0.122 | -0.520 |
|  | P | 9.11E-04 | 4.78E-01 | 5.07E-04 | 1.10E-01 | 4.95E-01 | 3.30E-03 | 3.40E-03 | 2.83E-01 | 3.40E-05 |
| BP-I | Estimate | -0.816 | -0.502 | -0.928 | -0.483 | -0.062 | -0.290 | -0.827 | -0.492 | -1.031 |
|  | P | 5.29E-18 | 8.30E-08 | 4.67E-20 | 5.64E-08 | 4.60E-01 | 5.42E-04 | 2.49E-17 | 3.31E-08 | 2.31E-25 |
| SCZ | Estimate | -1.976 | -1.262 | -1.256 | -0.515 | 0.331 | -0.167 | -1.697 | -0.499 | -1.318 |
|  | P | 1.83E-12 | 5.04E-06 | 1.98E-05 | 3.37E-02 | 1.76E-01 | 4.94E-01 | 3.59E-11 | 3.66E-02 | 4.49E-07 |
| Male | Estimate | 0.138 | 0.138 | 0.112 | 0.046 | 0.064 | -0.279 | -0.146 | -0.084 | 0.215 |
|  | P | 1.48E-01 | 1.39E-01 | 2.69E-01 | 6.13E-01 | 4.50E-01 | 7.79E-04 | 1.47E-01 | 3.60E-01 | 3.55E-02 |
| MDD:Male | Estimate | 0.156 | 0.256 | -0.137 | -0.074 | 0.044 | 0.323 | -0.096 | 0.037 | -0.122 |
|  | P | 2.65E-01 | 5.80E-02 | 3.54E-01 | 5.74E-01 | 7.22E-01 | 7.65E-03 | 5.07E-01 | 7.79E-01 | 4.08E-01 |
| BP-II:Male | Estimate | 0.329 | -0.004 | -0.045 | 0.157 | 0.076 | 0.314 | 0.076 | 0.046 | 0.262 |
|  | P | 1.36E-01 | 9.84E-01 | 8.49E-01 | 4.55E-01 | 6.97E-01 | 1.04E-01 | 7.41E-01 | 8.28E-01 | 2.60E-01 |
| BP-I:Male | Estimate | 0.108 | 0.257 | -0.111 | -0.033 | -0.006 | 0.204 | -0.076 | -0.010 | 0.090 |
|  | P | 4.69E-01 | 8.21E-02 | 4.87E-01 | 8.15E-01 | 9.65E-01 | 1.23E-01 | 6.23E-01 | 9.44E-01 | 5.66E-01 |
| SCZ:Male | Estimate | 1.212 | 0.718 | 0.088 | 0.053 | -0.486 | 0.188 | 0.406 | -0.083 | 0.145 |
|  | P | 9.43E-05 | 1.92E-02 | 7.87E-01 | 8.47E-01 | 7.30E-02 | 4.88E-01 | 1.63E-01 | 7.58E-01 | 6.25E-01 |

Additionally, there was a significant three-way interaction among sex, diagnosis and test domain for speed, indicating that the effect of sex on the speed of cognitive performance depended on diagnosis and test domain (Supplementary Table 5A). This interaction apparently resulted from the differentially slower performance of females with SCZ, BP-I, and MDD on attention and working memory tests (Supplementary Figure 3).

The largest deficits in cases, relative to controls, were seen in executive function speed and accuracy, especially attention and working memory; in social cognition, particularly emotion identification; and in motor speed. While most participants were taking medications (Table 1), conclusions in the above analyses were robust to inclusion of medication use as a covariate (Supplementary Table 5B).

### Symptom Endorsement in Cases

Twenty-one symptoms were endorsed by at least 2·5% of cases (Supplementary Figure 4). All symptoms were present in SCZ cases, whereas psychosis-associated symptoms were uncommon in BP-II and MDD cases. BP-I cases endorsed psychosis-associated symptoms at a reduced level compared to SCZ cases, however religious delusions were nearly as common in BP-I as in SCZ. Depressed mood, anhedonia, fatigue, avolition, and suicidal thoughts were common across all diagnoses.

### Associations among Symptom Factors and Cognitive Performance

To analyze the relationship between cognitive performance and psychiatric symptoms across diagnoses, we first performed a factor analysis of the binary symptom data from the 1,689 cases diagnosed with SCZ, BP-I, BP-II, and MDD; by doing so we could represent the 21 categorical, and collinear symptoms with prevalence in excess of 2·5% by a smaller number of continuous scores. This method has been applied previously to reduce dimensionality of symptom ratings (26). We determined that a three-factor model best represented the symptom data, as indicated by a combination of theoretical, empirical, and common subjective methods (22–24) (see Methods). The three symptom factors can be described as Psychosis (positive loadings for hallucinations, delusions, and disorganized speech/behavior), Mania (positive loadings for decreased need for sleep, flight of ideas and grandiosity, and negative loadings for avolition), and Depression (positive loadings for anhedonia, fatigue, depressed mood, hypersomnia, suicide attempt, and suicidal thoughts) (Supplementary Figure 5, Supplementary Table 6). As expected, SCZ cases are elevated on the Psychosis factor, BP-II and MDD cases are elevated on the Depression factor, and BP-I cases are elevated on the Mania factor; however, distributions of factor scores overlap substantially across diagnostic categories (Supplementary Figure 6).

Including only cases, we repeated the LMM analysis using factor scores on Psychosis, Mania, and Depression as covariates (Supplementary Table 5C). Controlling for diagnosis, high Psychosis factor score was significantly associated with reduced accuracy and slower speeds; high Depression score was significantly associated with increased accuracy; and Mania factor score was not significantly associated with accuracy or speed. While premorbid IQ (WAT) and both lifetime and current illness severity (number of hospital admissions/ER visits and SA-45 scores, respectively) were associated with both diagnosis and factor scores (Supplementary Table 7), conclusions in the above analyses were robust to inclusion of these data as covariates (Supplementary Tables 5D, 5E).

The three-way interaction among diagnosis, sex, and test domain was not significant for either speed or accuracy (Supplementary Table 5C); however, both speed and accuracy had significant two-way interactions of test domain with sex and with diagnosis. These interactions prompted us to analyze neurocognitive test domains individually, including main effects of diagnosis and sex. We included the factor scores that were significant in the combined LMM analyses as covariates in these analyses (Supplementary Table 5C, Supplementary Figure 2): analysis of accuracy included Psychosis and Depression, while analysis of speed included only Psychosis.

Higher Psychosis scores were specifically associated with lower accuracy and slower speed in both executive function and social cognition, and with slower motor speed, while higher Depression scores were specifically associated with improved social cognition accuracy (Table 3). To visualize the effect of the Psychosis and Depression factors on cognition, we first regressed the effects of diagnosis and sex out of raw CNB scores, prior to generating z-scores. We then categorized cases as above or below the median of each factor, irrespective of diagnosis (Table 1), transformed to z-scores based on the mean and SD in cases in the lower group, and plotted the cognition profiles for both groups in each factor (Figures 2, 3). After removing the effect of diagnosis, we see that cases in the upper half of the distribution of Psychosis factor scores have poorer performance on both speed and accuracy than do cases in the lower half of the distribution.

**Figure 2.**
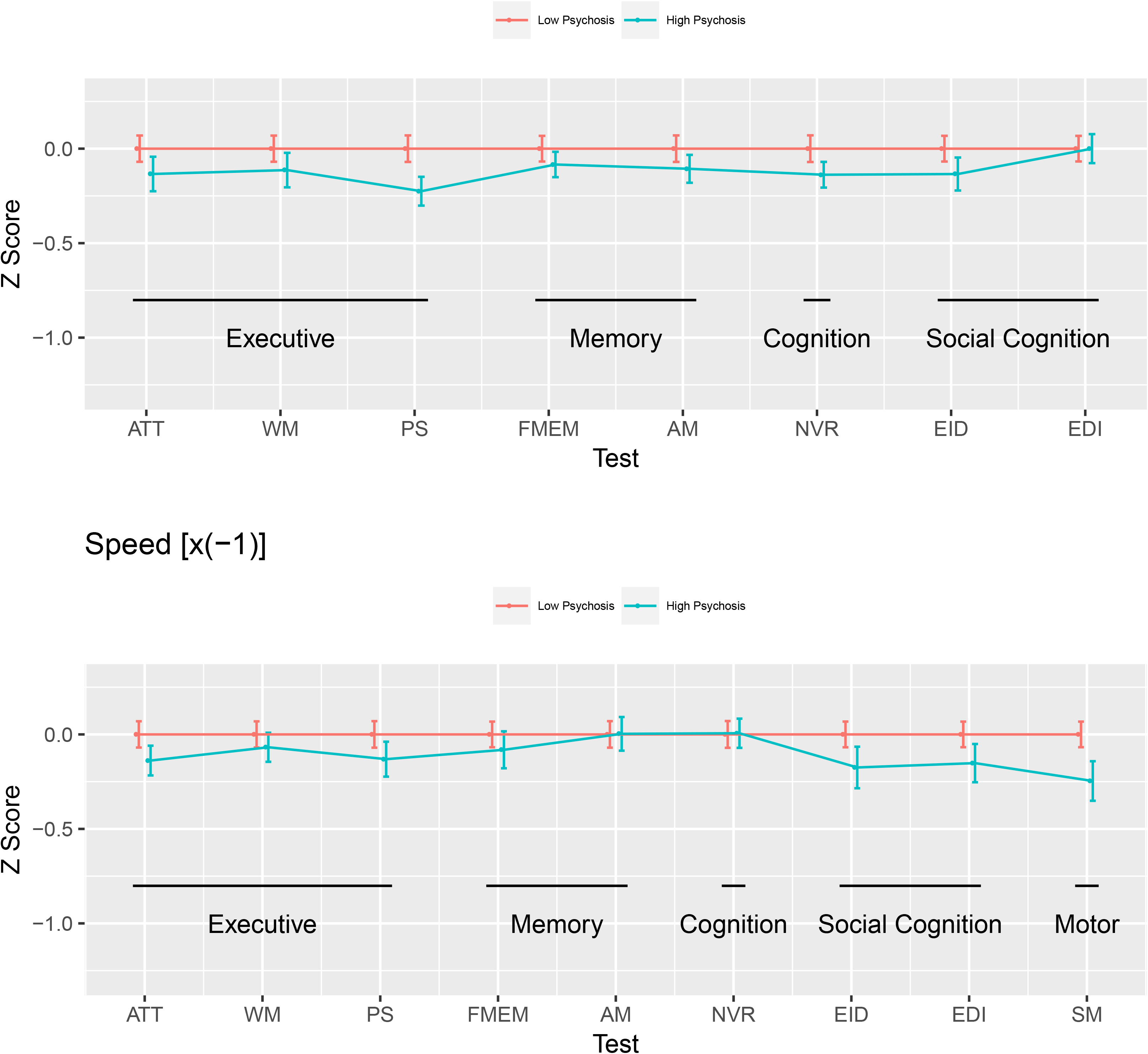
Z-scores on Accuracy (top) and Speed (bottom) profiles for tests assessing Executive Function, Memory, (Complex) Cognition, Social Cognition, and Motor Speed, stratified by Psychosis factor scores. Data for speed were multiplied by −1 so that poorer performance (slower speed), would result in a lower value. In order to focus on the effect of Psychosis factor score, diagnosis was regressed out of raw test data, and SMI cases were categorized as being above or below the median on the Psychosis factor score. Z-scores were generated relative to the low Psychosis group. Test abbreviations: ATT = Continuous Performance Test; WM = Letter-N-Back test; PS = Digit Symbol Test, matching trials; FMEM = Face Memory test; AM = Digit Symbol test, recall trials; NVR = Matrix Analysis test; EID = Emotion Identification test; EDI = Measured Emotion Differentiation test; SM = Motor Praxis test. High Psychosis=SMI cases with Psychosis factor scores above the median (n=831); Low Psychosis=SMI cases with Psychosis factor score below the median (n=858). Error bars are the 95% confidence intervals. The number of subjects in each Psychosis group, by diagnosis, is presented in Table 1.

**Figure 3.**
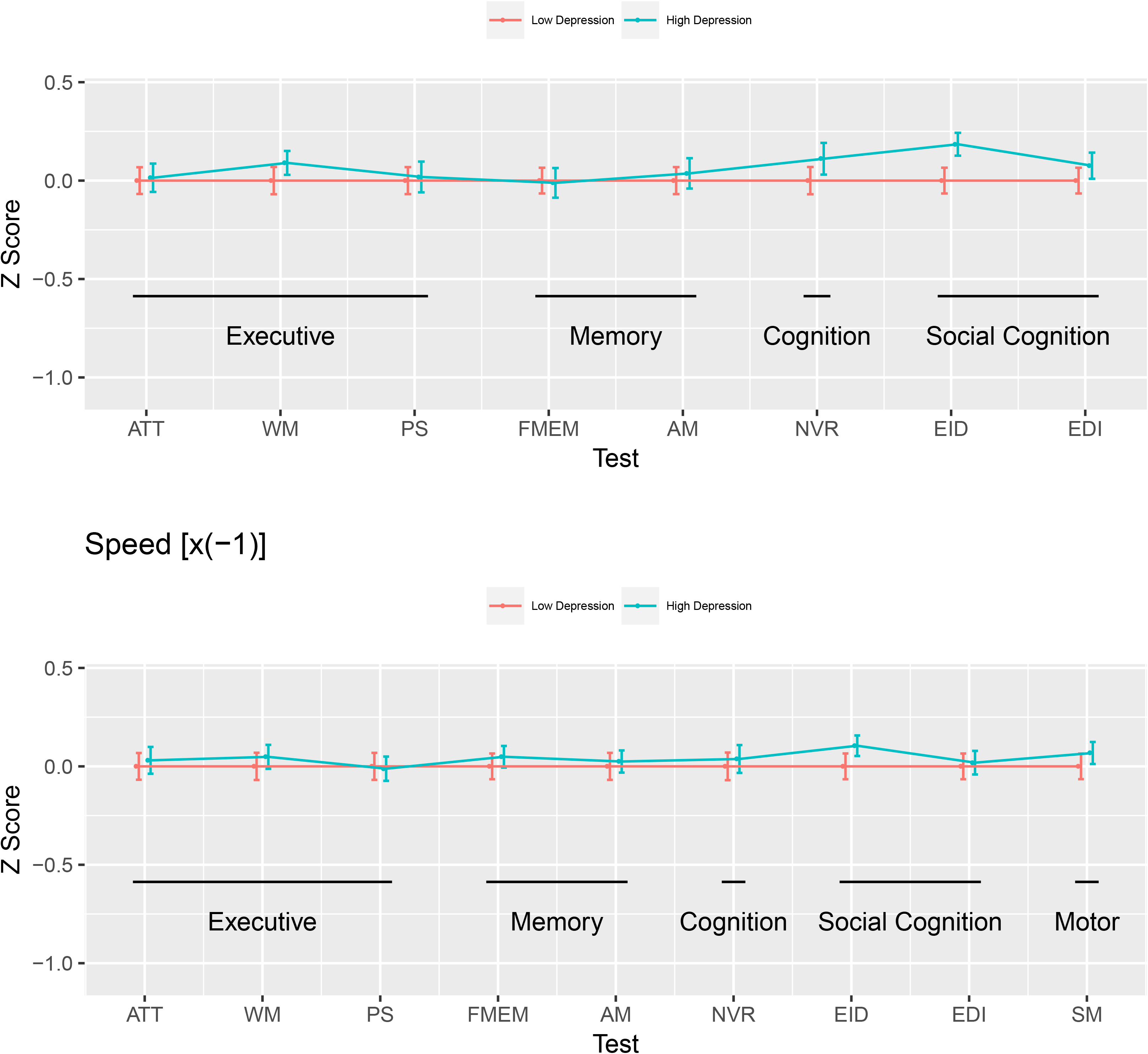
Z-scores on Accuracy (top) and Speed (bottom) profiles for tests assessing Executive Function, Memory, (Complex) Cognition, Social Cognition, and Motor Speed, stratified by Depression factor scores. Data for speed were multiplied by −1 so that poorer performance (slower speed), would result in a lower value. In order to focus on the effect of Depression factor score, diagnosis and sex were regressed out of raw test data, and SMI cases were categorized as being above or below the median on the Depression factor score. Z-scores were generated relative to the low Depression group. Test abbreviations: ATT = Continuous Performance Test; WM = Letter-N-Back test; PS = Digit Symbol Test, matching trials; FMEM = Face Memory test; AM = Digit Symbol test, recall trials; NVR = Matrix Analysis test; EID = Emotion Identification test; EDI = Measured Emotion Differentiation test; SM = Motor Praxis test. High Depression=SMI cases with Depression factor scores above the median (n=752); Low Depression=SMI cases with Depression factor score below the median (n=937). Error bars are the 95% confidence intervals. The number of subjects in each Depression group, by diagnosis, is presented in Table 1.

**Table 3.**
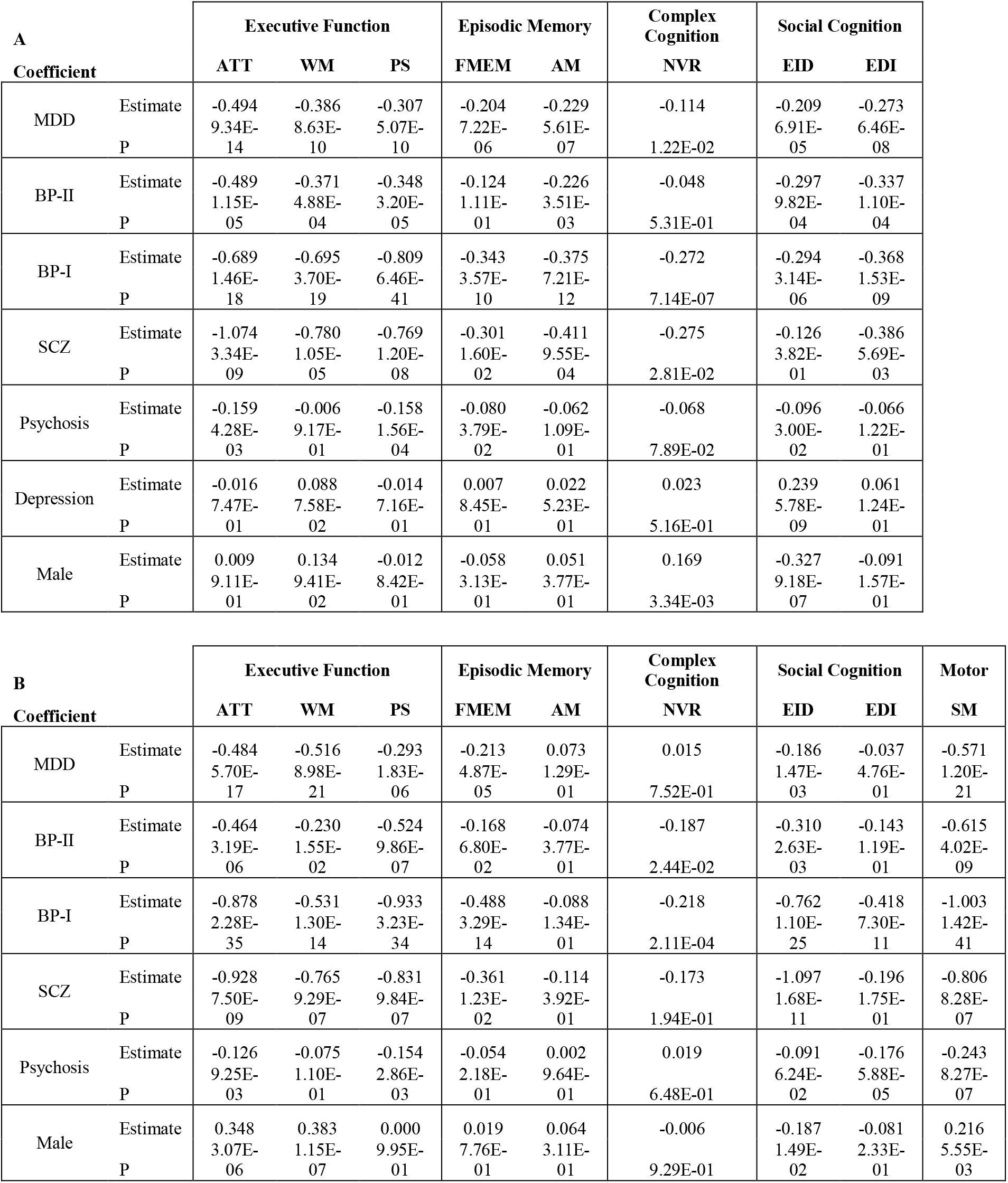
Analyses of accuracy (A) and speed (B) z-scores for each Penn-CNB test, including psychosis factor score as a covariate for speed, and psychosis and depression factor scores as covariates for accuracy. ATT = Continuous Performance Test; WM = Letter-N-Back test; PS = Digit Symbol Test, matching trials; FMEM = Face Memory test; AM = Digit Symbol test, recall trials; NVR = Matrix Analysis test; EID = Emotion Identification test; EDI = Measured Emotion Differentiation test; SM = Motor Praxis test.

In contrast, cases in the upper half of the distribution of Depression scores have improved social cognition accuracy.

## Discussion

In this study we investigated a prospective cohort that included large samples of cases representing each of the three main SMI diagnoses, SCZ, BPD, and MDD. The study design enables transdiagnostic analyses not possible in previous investigations, in that all cases were ascertained agnostic to diagnosis, and assessed uniformly regardless of diagnosis; both for performance across major neurocognitive domains and for individual lifetime symptoms. These assessments provide new insights on the magnitude and profile of neurocognitive impairment in SMI in relation to both diagnosis and empirically-derived, trans-diagnostic symptom factors. Additional assessments, including evaluation of EMRs available for most cases, allowed us to show that our findings were robust to lifetime and current illness severity, premorbid IQ, and medication usage.

The bifurcation of cognitive profiles was the most striking finding with respect to diagnoses: compared to healthy controls, SCZ and BP-I showed pronounced deficits across executive function, social cognition and motor speed tests relative to memory and complex cognition, while BP-II and MDD displayed mild cognitive impairment, across domains, with intact nonverbal reasoning. This result aligns with a growing body of evidence indicating heterogeneity of neurocognitive function within BPD (27), and provides a possible explanation for a similarly bifurcated pattern of genetic correlations between the SMI diagnoses revealed in recent large-scale GWAS datasets (28). The large trans-diagnostic sample also enabled us to identify a significant interaction between test domain, diagnosis and sex, as females with SCZ, BP-I, and MDD were slower than males in measures of executive speed.

Because of the branching structure of diagnostic interviews, SMI symptoms are usually assessed only in respondents who endorse screening questions for the diagnoses typically associated with those symptoms. By uniformly assessing lifetime symptomatology outside of the NetSCID, we found that a substantial number of symptoms (depressed mood, anhedonia, fatigue, avolition, and suicidal thoughts) occurred at a relatively high frequency (> 25%) across all diagnoses. Our study design is ideal for further investigation of such symptoms, for example, through analyses aimed at dissecting polygenic risk across SMI diagnoses. At that same time, further explorations of the symptom data may shed light on the biology related to specific diagnoses. While the lifetime frequencies of the overall set of psychosis symptoms aligned with the above-noted bifurcation of cognition profiles (high in SCZ/BP-I and low in BP-II/MDD), this pattern reflects mainly a few symptoms that are nearly as prominent in BP-I as in SCZ (e.g. religious delusions).

The exploratory item-factor analysis of clinical symptoms identified a clear psychosis factor and two mood-related factors, which enabled trans-diagnostic evaluations of symptomatology in relation to specific neurocognitive domains. Even after adjusting for effects of diagnosis, increasing scores on the Psychosis factor were associated with slower motor speed and reduced accuracy in both executive function and social cognition. Previous studies have shown similar associations (29), but have not examined such a broad range of SMI diagnoses, or such a uniformly ascertained study population. We also obtained the unexpected finding that higher scores on the Depression factor were associated with improved social cognition accuracy, after adjusting for effects of diagnosis; our results suggest that the effect, cross-diagnostically, is larger for accuracy rather than speed. For both symptom factors, effects persisted after controlling for disease severity and premorbid IQ.

It is now well-accepted that premorbid cognitive impairment is a common feature of SCZ (30). While we found that the effects of the Psychosis symptom factor persisted after controlling for premorbid IQ, our results cannot differentiate between two possible mechanisms: (a) lifetime psychotic symptomatology has deleterious effects on specific cognitive domains and these effects transcend diagnostic categories; or (b) a common set of risk factors may predispose to both psychotic symptoms and impaired cognition across SMI categories. Longitudinal data, from a developmental perspective, could shed light on this mechanism; further investigation of the EMRs of the Paisa cohort may provide such information.

We note two limitations of this study. First, although SMI case and control participants were recruited from the same communities, we cannot rule out subtle effects of demographic differences between these groups. We adjusted for any such effects statistically. Second, while previous studies have highlighted deficits in verbal memory in SCZ (30), we did not assess this domain due to lack of normative data on word frequency in the Paisa population.

Independent studies have shown that the specific cognitive domains measured here are heritable (31), while studies of more limited sets of cognitive measures have demonstrated their genetic correlation with SCZ (32, 33). Future genome-wide genotyping studies of the Paisa cohort described here will enable the examination of genetic correlations for neurocognitive measures from the current study across SMI diagnoses and symptom factors, broadly, as well as GWAS of neurocognitive phenotypes across multiple domains. This large cohort of uniformly ascertained individuals thus provides unique opportunities for the phenotypic and genetic characterization of SMI that may ultimately lead to novel approaches for disease classification.

## Acknowledgments

This study was supported by NIMH grants R01MH113078 (to NBF, CEB, and CLJ); R01MH095454-02S1 to NBF; and K99MH116115 to LOL. CTSI Grant UL1TR001881 provided support for the data collection system REDCap. The authors thank the CSJDM and HUSVF and all study participants for making this work possible.

## Declaration of Interest

Dr. Benjamin Brodey owns stock in TeleSage, Inc. TeleSage, Inc. created the NetSCID software using funding from the NIH and licenses the SCID from the American Psychiatric Press. All other authors have no declarations.

## Author Contributions

CVU, MCR, LGAA, AMDZ, JME, MCL, JDP, SSR, JV recruited and interviewed participants and administered test instruments. TT, MMU, and AE designed data bases, coordinated data transfers and managed data downloads. LOL and JDHG provided EMR data. BB modified NetSCID for our use. CS, SKS, and TMM were responsible for data analysis. JIE, VIR, CLJ, RCG, CEB, and NBF designed the study. SKS, CEB, RCG, and NBF wrote the paper.

## Notes

#### Summary of Updates

Increased sample size, additional analyses of severity, Supplemental files updated

