## Supplementary Figures and Tables for "Distinct and shared contributions of diagnosis and symptom domains to cognitive performance in a case-control study of severe mental illness in the Paisa population"

**Supplementary Figure 1. Flow chart of participant recruitment.** Incomplete assessment: enrolled, but interview/assessments not yet finished. No CNB: participants were unable to complete CNB due to fatigue, problems with computer use, or mental state (see Supplementary Table 4 for details). Invalid CNB: CNB was attempted, but participant was unable to do assessment without help. Non-target DX: case participants with DX other than SCZ, BP-I, BP-II, or MDD

**Supplementary Figure 2. Schematic of linear mixed model analysis procedures and overview of results.**

**Supplementary Figure 3. Mean Speed z-scores stratified by test domain, diagnosis and sex. Z-scores were generated relative to Controls.** Error bars are the 95% confidence intervals. Data for Speed were multiplied by -1 so that poorer performance (slower speed), would result in a lower value. F=female, M=male. Test abbreviations: ATT = Continuous Performance Test; WM = Letter-N-Back test; PS = Digit Symbol Test, matching trials; FMEM = Face Memory test; AM = Digit Symbol test, recall trials; NVR = Matrix Analysis test; EID = Emotion Identification test; EDI = Measured Emotion Differentiation test; SM = Motor Praxis test.

**Supplementary Figure 4. Symptom endorsement by diagnosis for 21 binary symptoms endorsed by at least 2.5% of cases overall.** SCZ: schizophrenia (n=160) BP-I: bipolar disorder I (n=519), BP-II: bipolar disorder II (n=204), MDD: major depressive disorder (n=806).

**Supplementary Figure 5. Factor loadings on 21 binary symptoms in the three-factor model.**

**Supplementary Figure 6. Distribution of scores on factors, by diagnosis.** SCZ: schizophrenia (n=160) BP-I: bipolar disorder I (n=519), BP-II: bipolar disorder II (n=204), MDD: major depressive disorder (n=806).

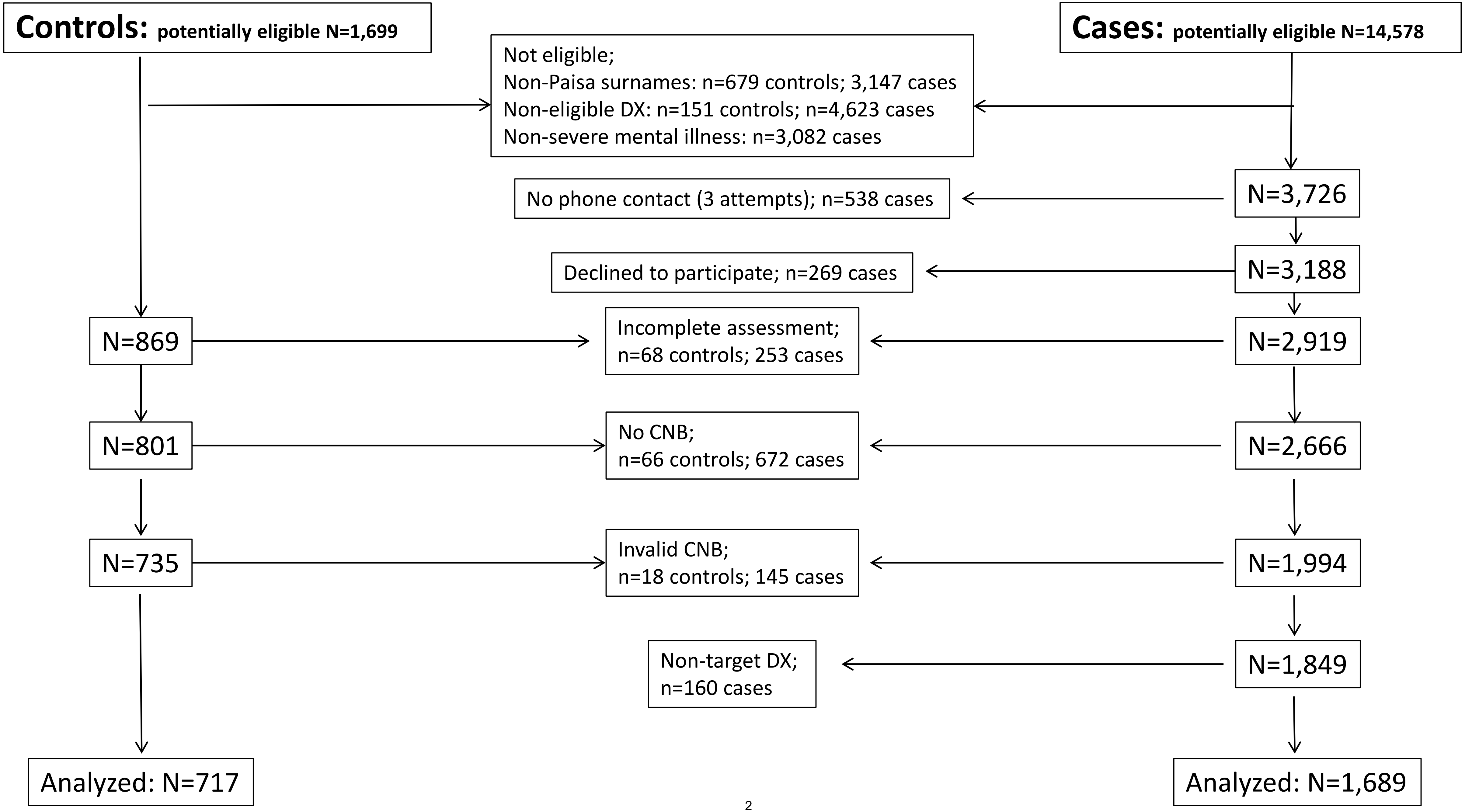
Controls: potentially eligible N=1,699
Cases: potentially eligible N=14,578
Not eligible;
Non-Paisa surnames: n=679 controls; 3,147 cases
Non-eligible DX: n=151 controls; n=4,623 cases
Non-severe mental illness: n=3,082 cases
N=3,726
No phone contact (3 attempts); n=538 cases
N=3,188
Declined to participate; n=269 cases
N=869
Incomplete assessment;
n=68 controls; 253 cases
N=2,919
N=801
No CNB;
n=66 controls; 672 cases
N=2,666
N=735
Invalid CNB;
n=18 controls; 145 cases
N=1,994
Non-target DX;
n=160 cases
N=1,849
Analyzed: N=717
Analyzed: N=1,689

Supplementary Figure 2

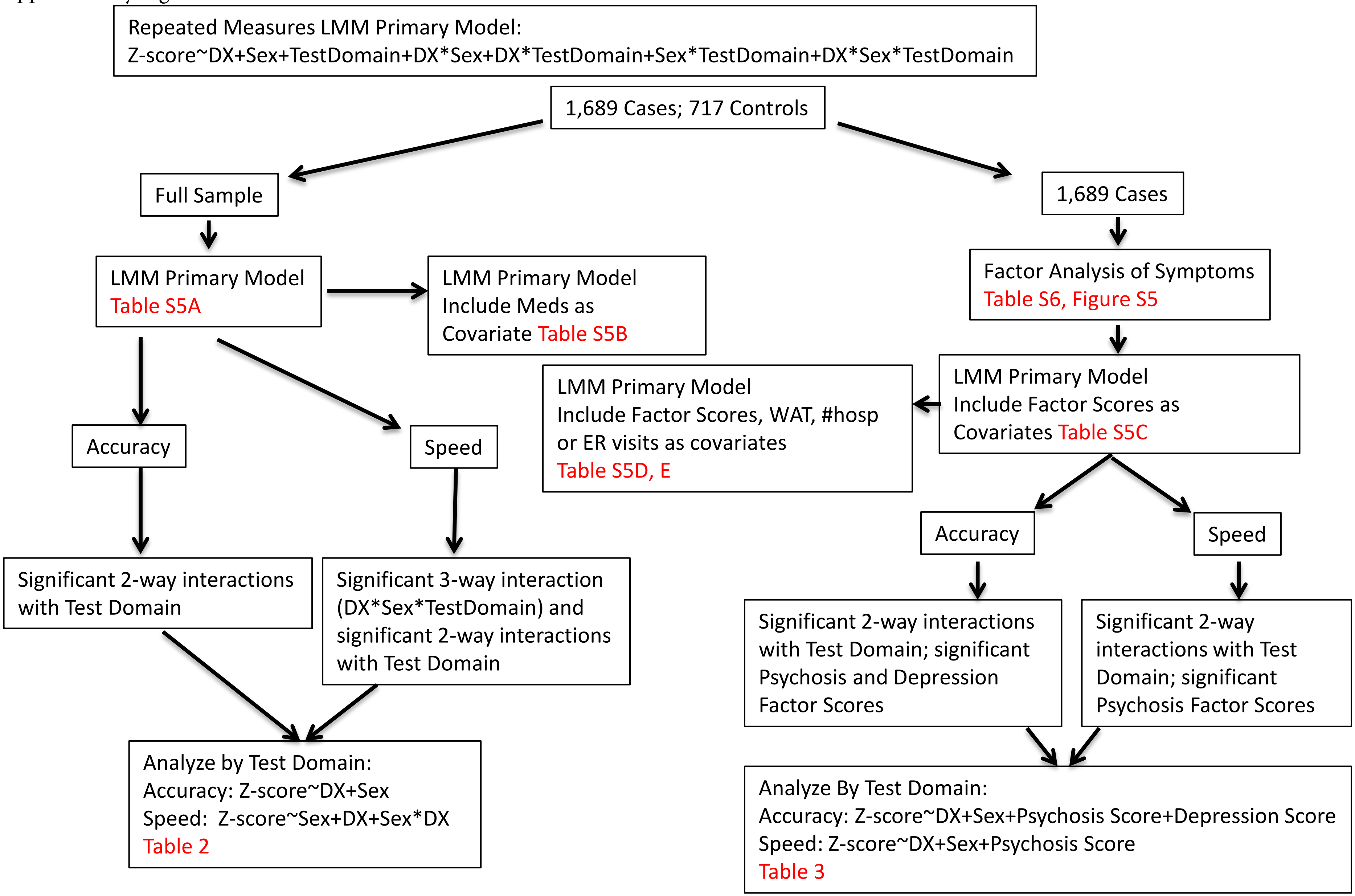

Supplementary Figure 3

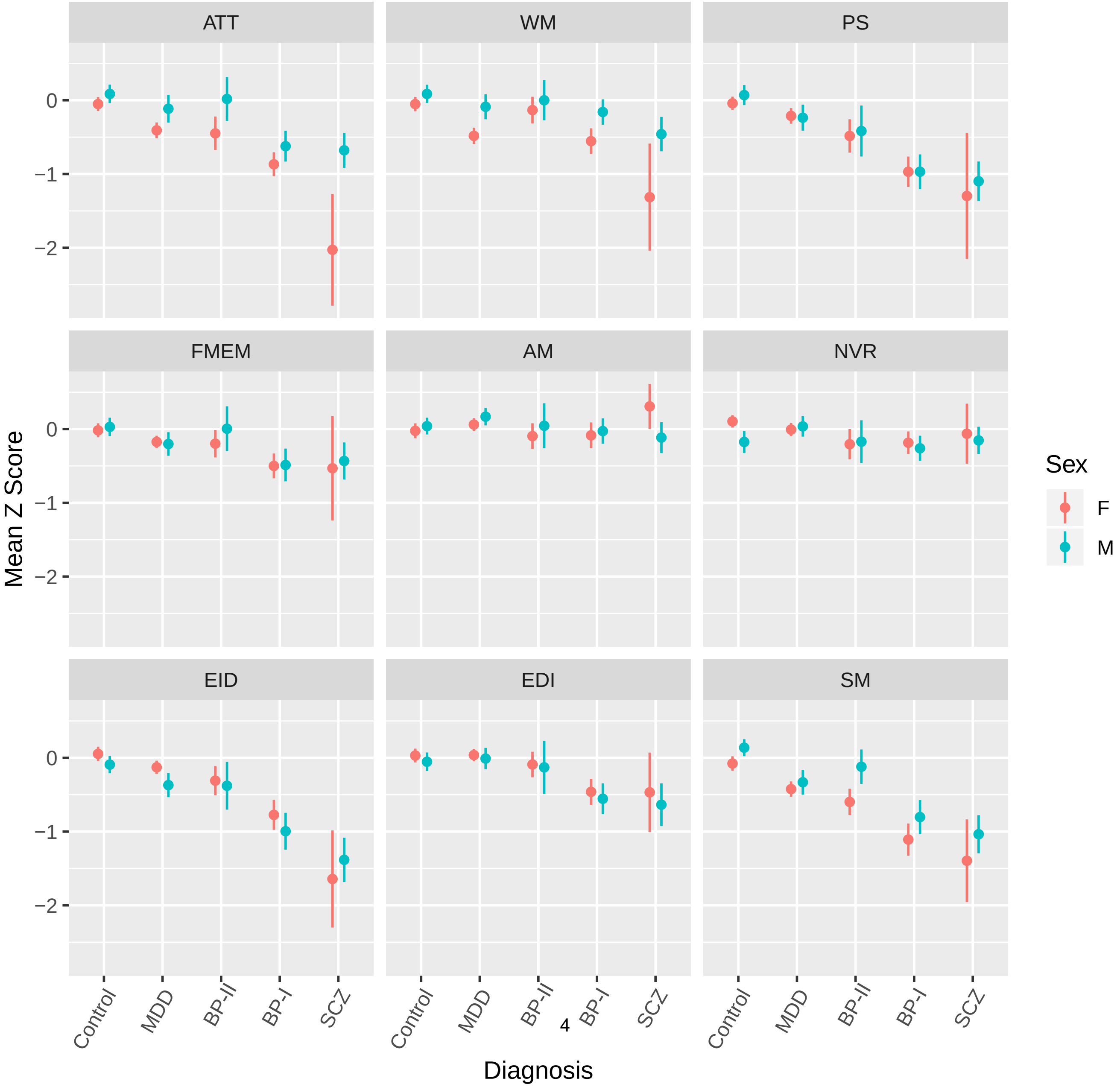

Supplementary Figure 4

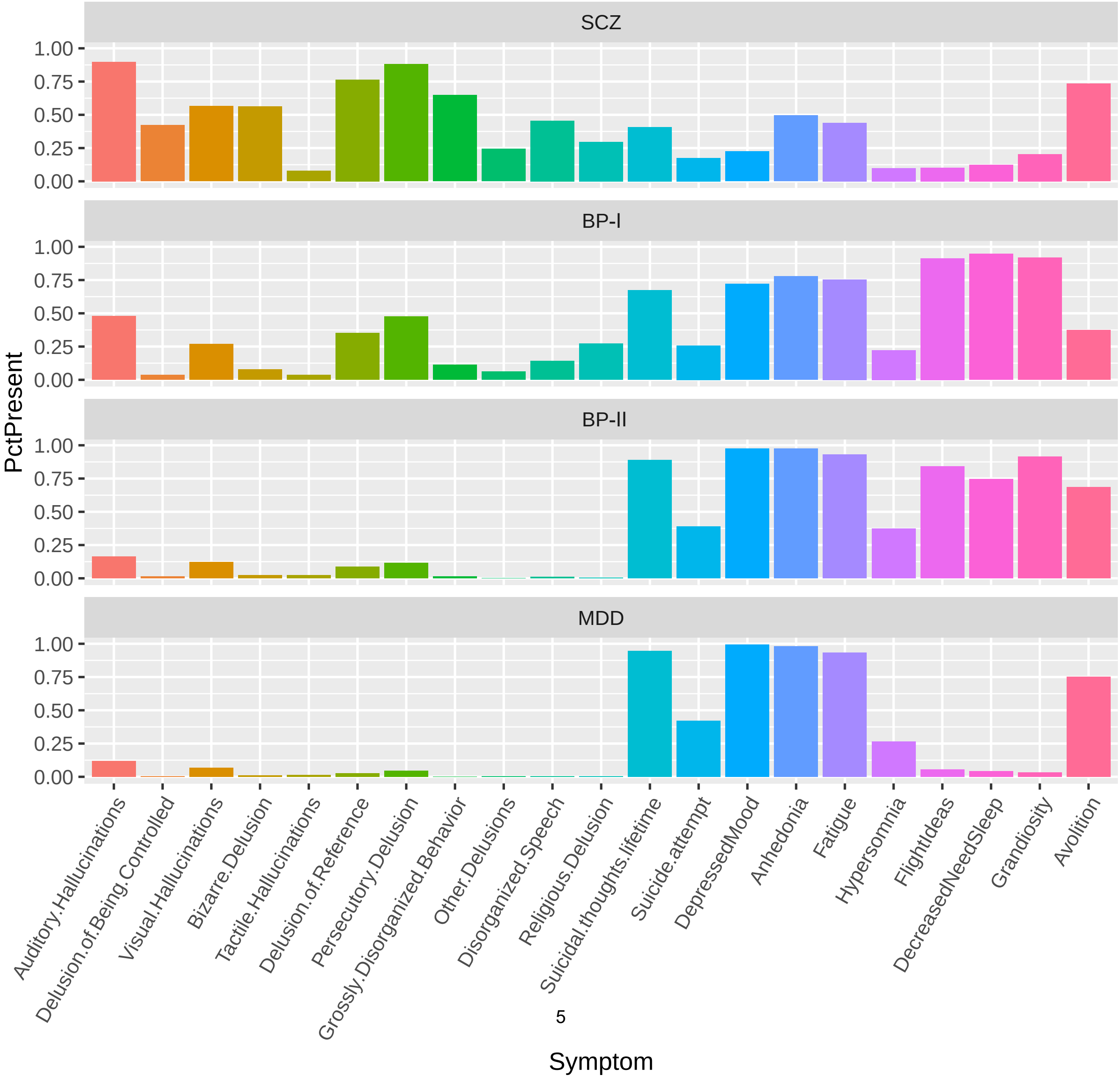

Supplementary Figure 5

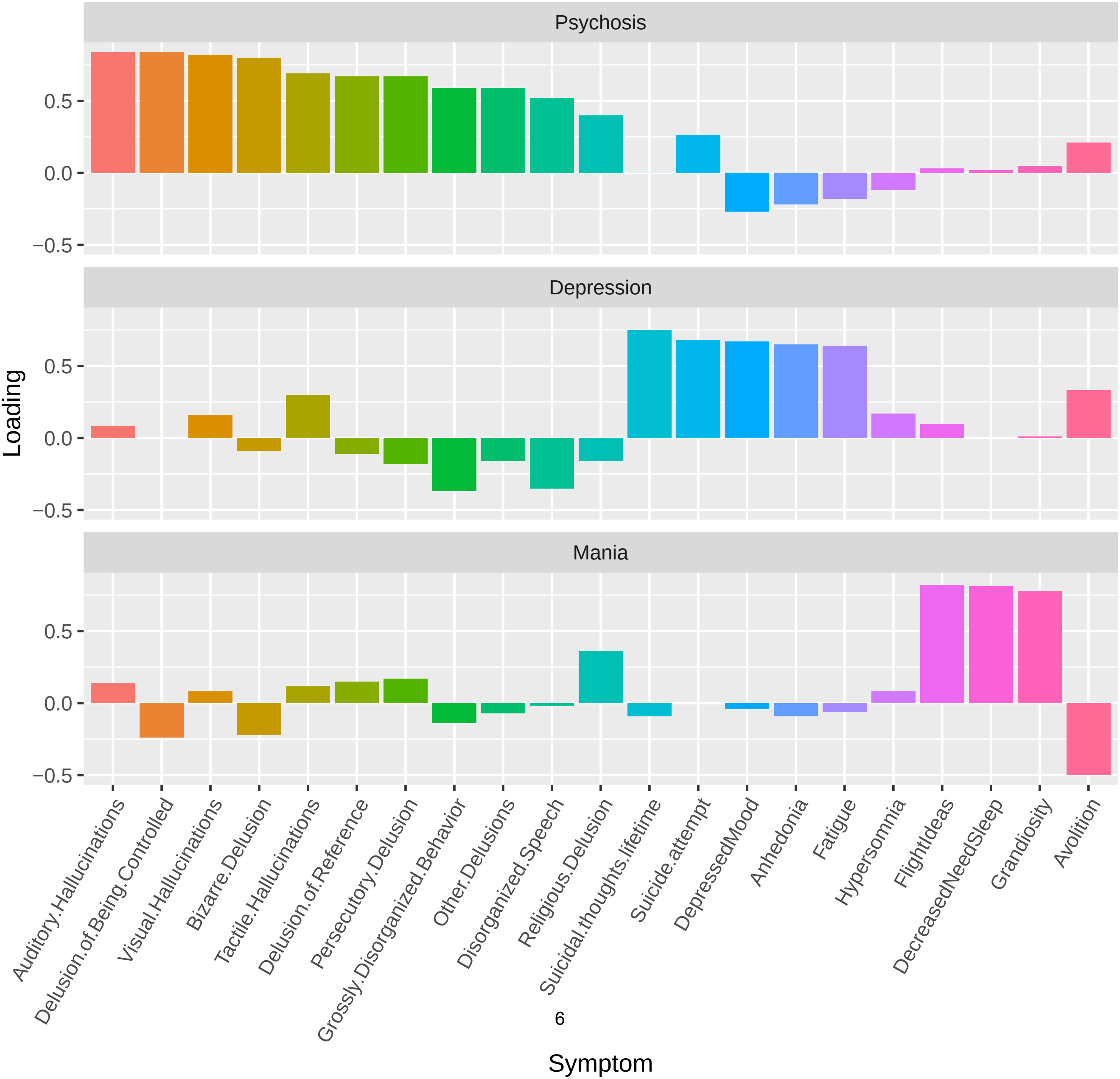

Supplementary Figure 6

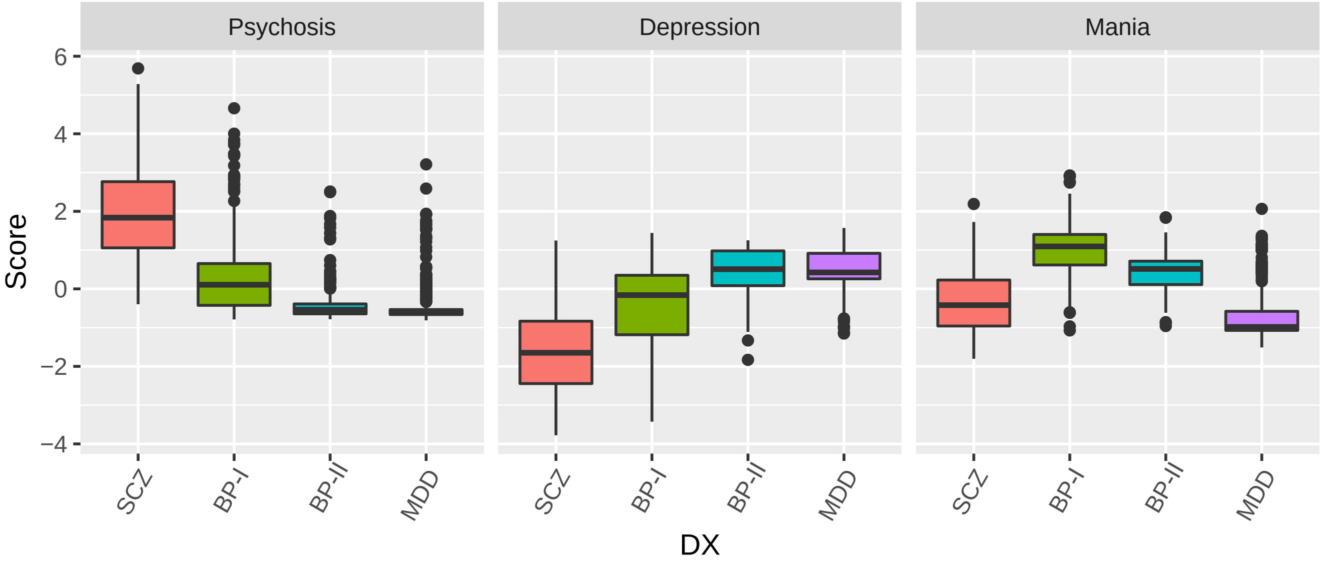

**Supplementary Table 1. Summary of missing data for variables analysed in 1,689 cases and 717 controls**

| Category | Variable | Quantitative or Categorical | Outcome or predictor | N Missing Case | N Missing Control |
|---|---|---|---|---|---|
| Penn CNB Speed; see Supplementary Table 1 for detail | SM | Quantitative | Outcome | 41 | 8 |
| | EDI | | | 51 | 9 |
| | EID | | | 71 | 13 |
| | WM | | | 195 | 34 |
| | FMEM | | | 61 | 13 |
| | AM | | | 194 | 27 |
| | PS | | | 194 | 27 |
| | ATT | | | 163 | 17 |
| | NVR | | | 247 | 42 |
| Penn CNB Accuracy see Supplementary Table 1 for detail | EDI | Quantitative | Outcome | 51 | 9 |
| | EID | | | 71 | 13 |
| | WM | | | 194 | 34 |
| | FMEM | | | 61 | 13 |
| | AM | | | 194 | 27 |
| | PS | | | 194 | 27 |
| | ATT | | | 162 | 16 |
| | NVR | | | 218 | 35 |
| Symptom | Auditory Hallucinations | Categorical | Predictor | 15 | NA |
| | Delusion of Being Controlled | | | 30 | |
| | Visual Hallucinations | | | 34 | |
| | Bizarre Delusion | | | 27 | |
| | Tactile Hallucinations | | | 28 | |
| | Delusion of Reference | | | 32 | |
| | Persecutory Delusion | | | 13 | |
| | Grossly Disorganized Behavior | | | 16 | |
| | Other Delusions | | | 33 | |
| | Disorganized Speech | | | 14 | |
| | Religious Delusion | | | 23 | |
| | Suicidal thoughts lifetime | | | 24 | |
| | Suicide attempt | | | 22 | |
| | Depressed Mood | | | 1 | |
| | Anhedonia | | | 74 | |
| | Fatigue | | | 127 | |
| | Hypersomnia | | | 127 | |
| | Flight of Ideas | | | 214 | |
| | Decreased Need for Sleep | | | 207 | |
| | Grandiosity | | | 96 | |
| | Avolition | | | 2 | |
| Factor Scores | Psychosis | Quantitative | Predictor | 0 | NA |
| | Depression | | | 0 | |
| | Mania | | | 0 | |
| pre-morbid IQ | WAT | Quantitative | Predictor | 7 | 2 |
| disease severity | number of hospitalizations and ER visits | Quantitative | Predictor | 359 | NA |
| current symptom severity | SA45 | Quantitative | Predictor | 90 | 46 |

**Supplementary Table 2. Description of tests employed in the Penn CNB**

| Test | Description | Abbreviation | RDoC Domain |
|---|---|---|---|
| Motor Praxis | 40 total trials, including 20 test trials, to measure ability to control a computer mouse | **SM** | Sensorimotor |
| Measured Emotion Intensity Differentiation | 36 paired faces varying in intensity to evaluate participant's ability for intensity differentiation as part of social cognition function | **EDI** | Social Communication |
| Penn Emotion Identification | 40 faces presented with emotions that range from mild to extreme intensity that participants have to identify to evaluate social cognition function | **EID** | |
| Short Letter-N-Back | 90 total stimuli to assess participant's working memory as part of executive functioning | **WM** | Working Memory |
| Penn Face Memory | 20 faces to memorize one at a time to assess episodic memory function | **FMEM** | Declarative Memory |
| Digit Symbol | numbers paired with symbols to assess participant's speed of information processing domain (Processing Speed) as well as incidental learning (Associative Memory) | **AM** | |
| | | **PS** | Attention |
| Short Penn Continuous Performance | 180 total trials including 90 number and 90 letter trials to assess participant's Attention as part of executive function | **ATT** | |
| Penn Matrix Analysis | 24 items with increasing difficulty to evaluate participant's complex cognition | **NVR** | Global IQ |

**Supplementary Table 3. Cases were asked about their current use of the following medications.** They were grouped into Antidepressants, Antipsychotics, and Mood Stabilizers for analysis; a 1 in a column indicates the group to which the medication belongs. N=number of patients who report currently taking the medication

| Drug | N | Antidepressant | Antipsychotic | Mood Stabilizer |
|---|---|---|---|---|
| Valproic Acid | 487 | 0 | 0 | 1 |
| Agomelatine | 55 | 1 | 0 | 0 |
| Amisulpride | 39 | 0 | 1 | 0 |
| Amitriptyline | 18 | 1 | 0 | 0 |
| Aripiprazole | 40 | 0 | 1 | 0 |
| Asenapine | 1 | 0 | 1 | 0 |
| Bupropion | 33 | 1 | 0 | 0 |
| Carbamazepine | 20 | 0 | 0 | 1 |
| Lithium Carbonate | 175 | 0 | 0 | 1 |
| Clomipramine | 2 | 1 | 0 | 0 |
| Clozapine | 135 | 0 | 1 | 0 |
| Desvenlafaxine | 11 | 1 | 0 | 0 |
| Duloxetine | 29 | 1 | 0 | 0 |
| Escitalopram | 204 | 1 | 0 | 0 |
| Fluoxetine | 146 | 1 | 0 | 0 |
| Fluvoxamine | 36 | 1 | 0 | 0 |
| Haloperidol | 8 | 0 | 1 | 0 |
| Lamotrigine | 38 | 0 | 0 | 1 |
| Levomepromazine | 68 | 0 | 1 | 0 |
| Mirtazapine | 9 | 1 | 0 | 0 |
| Olanzapine | 85 | 0 | 1 | 0 |
| Paliperidone | 3 | 0 | 1 | 0 |
| Paroxetine | 15 | 1 | 0 | 0 |
| Pipotiazine | 8 | 0 | 1 | 0 |
| Quetiapine | 447 | 0 | 1 | 0 |
| Risperidone | 180 | 0 | 1 | 0 |
| Sertraline | 263 | 1 | 0 | 0 |
| Trazodone* | 184 | 1 | 0 | 0 |
| Venlafaxine | 39 | 1 | 0 | 0 |
| Vortioxetine | 2 | 1 | 0 | 0 |

*only doses in excess of 300mg were classified as antidepressants

**Supplementary Table 4. Breakdown of reasons for missing CNB data.**

| Reason | N Cases | % Cases Missing Data | N Controls | % Controls Missing Data |
|---|---|---|---|---|
| Invalid CNB* | 145 | 17.75% | 18 | 21.43% |
| Lack of computer skills | 194 | 23.75% | 23 | 27.38% |
| Physical limitations** | 130 | 15.91% | 11 | 13.10% |
| Symptomatic | 105 | 12.85% | 0 | 0.00% |
| Illiterate | 45 | 5.51% | 4 | 4.76% |
| Cognitive Impairment | 33 | 4.04% | 0 | 0.00% |
| Fatigue | 4 | 0.49% | 0 | 0.00% |
| Declined to participate | 46 | 5.63% | 8 | 9.52% |
| Technical or Logistical problems in test administration*** | 75 | 9.18% | 16 | 19.05% |
| Unknown | 40 | 4.90% | 4 | 4.76% |
| **Total** | **817** | **100%** | **84** | **100%** |

*Testing was initiated and halted before completion when assessors realized participants were either symptomatic, or lacked computer skills needed to complete the assessment

**Physical limitations included, but were not limited to, osteoarthritis, fibromyalgia, tremor, hand surgery, carpal tunnel, visual limitations

***Problems include internet connectivity problems and lack of time

**Supplementary Table 5A: Analysis of accuracy and speed as a function of diagnosis, sex, domain, and interactions among main effects.** The dependent variable in analyses are z-scores on either Accuracy or Speed, the different tests administered are repeated measures in the LMM. Reference category is control-females. DX=Diagnosis. Prior to analysis, CNB data were adjusted for age and education.

| | Accuracy LMM N=2,406 | | | | Speed LMM N=2,406 | | | |
|---|---|---|---|---|---|---|---|---|
| | numDF | denDF | F-value | p-value | numDF | denDF | F-value | p-value |
| (Intercept) | 1 | 15453 | 394.5662 | <.0001 | 1 | 17761 | 300.4544 | <.0001 |
| DX | 4 | 2396 | 78.8939 | <.0001 | 4 | 2396 | 68.28848 | <.0001 |
| Sex | 1 | 2396 | 1.4032 | 0.2363 | 1 | 2396 | 4.70096 | 0.0302 |
| Domain | 7 | 15453 | 26.2324 | <.0001 | 8 | 17761 | 50.46407 | <.0001 |
| DX:Sex | 4 | 2396 | 2.1516 | 0.0721 | 4 | 2396 | 0.80256 | 0.5234 |
| DX:Domain | 28 | 15453 | 6.322 | <.0001 | 32 | 17761 | 13.19777 | <.0001 |
| Sex:Domain | 7 | 15453 | 9.937 | <.0001 | 8 | 17761 | 13.54875 | <.0001 |
| DX:Sex:Domain | 28 | 15453 | 1.1536 | 0.2627 | 32 | 17761 | 1.705 | 0.0078 |

**Supplementary Table 5B: Analysis of accuracy and speed as a function of diagnosis, sex, domain, interactions among these main effects, and medication use.** The dependent variable in analyses are z-scores on either Accuracy or Speed, the different tests administered are repeated measures in the LMM. Reference category is control-females. DX=Diagnosis. Prior to analysis, CNB data were adjusted for age and education.

| | Accuracy LMM N=2,406 | | | | Speed LMM N=2,406 | | | |
|---|---|---|---|---|---|---|---|---|
| | numDF | denDF | F-value | p-value | numDF | denDF | F-value | p-value |
| (Intercept) | 1 | 15453 | 400.12 | <.0001 | 1 | 17761 | 302.79 | <.0001 |
| DX | 4 | 2393 | 80.06 | <.0001 | 4 | 2393 | 68.84 | <.0001 |
| Sex | 1 | 2393 | 1.43 | 0.2326 | 1 | 2393 | 4.74 | 0.0296 |
| Domain | 7 | 15453 | 26.23 | <.0001 | 8 | 17761 | 50.48 | <.0001 |
| On Antipsychotics | 1 | 2393 | 26.41 | <.0001 | 1 | 2393 | 6.95 | 0.0084 |
| On Antidepressants | 1 | 2393 | 3.96 | 0.0466 | 1 | 2393 | 9.94 | 0.0016 |
| On Mood Stabilizers | 1 | 2393 | 9.79 | 0.0018 | 1 | 2393 | 7.74 | 0.0054 |
| DX:Sex | 4 | 2393 | 1.94 | 0.1013 | 4 | 2393 | 0.83 | 0.5089 |
| DX:Domain | 28 | 15453 | 6.32 | <.0001 | 32 | 17761 | 13.19 | <.0001 |
| Sex:Domain | 7 | 15453 | 9.93 | <.0001 | 8 | 17761 | 13.53 | <.0001 |
| DX:Sex:Domain | 28 | 15453 | 1.14 | 0.2734 | 32 | 17761 | 1.70 | 0.0079 |

**Supplementary Table 5C: Analysis of accuracy and speed as a function of diagnosis, sex, domain, interactions among these main effects, and scores on three symptom factors.** The dependent variable in analyses are z-scores on either Accuracy or Speed, the different tests administered are repeated measures in the LMM. DX=Diagnosis. Prior to analysis, CNB data were adjusted for age and education.

| | Accuracy LMM N=1,689 | | | | Speed LMM N=1,689 | | | |
|---|---|---|---|---|---|---|---|---|
| | numDF | denDF | F-value | p-value | numDF | denDF | F-value | p-value |
| DX | 4 | 1678 | 157.90217 | <.0001 | 4 | 1678 | 123.57637 | <.0001 |
| Sex | 1 | 1678 | 0.33674 | 0.5618 | 1 | 1678 | 5.01489 | 0.0253 |
| Domain | 7 | 10623 | 35.47571 | <.0001 | 8 | 12232 | 66.37686 | <.0001 |
| Psychosis | 1 | 1678 | 20.90826 | <.0001 | 1 | 1678 | 19.917 | <.0001 |
| Mania | 1 | 1678 | 4.65409 | 0.0311 | 1 | 1678 | 1.31302 | 0.252 |
| Depression | 1 | 1678 | 10.54576 | 0.0012 | 1 | 1678 | 2.22876 | 0.1357 |
| DX:Sex | 3 | 1678 | 0.31288 | 0.8161 | 3 | 1678 | 0.66007 | 0.5766 |
| DX:Domain | 21 | 10623 | 4.16805 | <.0001 | 24 | 12232 | 8.63453 | <.0001 |
| Sex:Domain | 7 | 10623 | 7.54731 | <.0001 | 8 | 12232 | 10.11055 | <.0001 |
| DX:Sex:Domain | 21 | 10623 | 1.05686 | 0.3886 | 24 | 12232 | 1.39371 | 0.0953 |

**Supplementary Table 5D: Analysis of accuracy and speed as a function of diagnosis, sex, domain, interactions among these main effects, scores on three symptom factors and score on the WAT.** The dependent variable in analyses are z-scores on either Accuracy or Speed, the different tests administered are repeated measures in the LMM. DX=Diagnosis. Prior to analysis, CNB data were adjusted for age.

| | Accuracy LMM N=1,682 | | | | Speed LMM N=1,682 | | | |
|---|---|---|---|---|---|---|---|---|
| | numDF | denDF | F-value | p-value | numDF | denDF | F-value | p-value |
| DX | 4 | 1670 | 202.35 | <.0001 | 4 | 1670 | 114.68 | <.0001 |
| Sex | 1 | 1670 | 0.00 | 0.9938 | 1 | 1670 | 9.00 | 0.0027 |
| Domain | 7 | 10595 | 32.26 | <.0001 | 8 | 12197 | 48.43 | <.0001 |
| wat_total | 1 | 1670 | 856.92 | <.0001 | 1 | 1670 | 334.03 | <.0001 |
| Psychosis | 1 | 1670 | 13.89 | 0.0002 | 1 | 1670 | 15.33 | 0.0001 |
| Mania | 1 | 1670 | 0.54 | 0.4627 | 1 | 1670 | 4.66 | 0.031 |
| Depression | 1 | 1670 | 4.85 | 0.0278 | 1 | 1670 | 0.52 | 0.47 |
| DX:Sex | 3 | 1670 | 0.31 | 0.8195 | 3 | 1670 | 1.42 | 0.234 |
| DX:Domain | 21 | 10595 | 4.19 | <.0001 | 24 | 12197 | 9.88 | <.0001 |
| Sex:Domain | 7 | 10595 | 7.55 | <.0001 | 8 | 12197 | 9.31 | <.0001 |
| DX:Sex:Domain | 21 | 10595 | 1.03 | 0.4214 | 24 | 12197 | 1.46 | 0.0669 |

**Supplementary Table 5E: Analysis of accuracy and speed as a function of diagnosis, sex, domain, interactions among these main effects, scores on three symptom factors, current symptom severity from the SA45, and number of hospitalizations and ER visits recorded in the electronic medical record.** The dependent variable in analyses are z-scores on either Accuracy or Speed, the different tests administered are repeated measures in the LMM. DX=Diagnosis. Prior to analysis, CNB data were adjusted for age and education.

| | Accuracy LMM N=1,330 | | | | Speed LMM N=1,330 | | | |
|---|---|---|---|---|---|---|---|---|
| | numDF | denDF | F-value | p-value | numDF | denDF | F-value | p-value |
| DX | 4 | 1251 | 137.42 | <.0001 | 4 | 1251 | 90.73 | <.0001 |
| Sex | 1 | 1251 | 0.28 | 0.5941 | 1 | 1251 | 3.93 | 0.0477 |
| Domain | 7 | 7762 | 27.25 | <.0001 | 8 | 8957 | 50.81 | <.0001 |
| Number of visits | 1 | 1251 | 0.80 | 0.371 | 1 | 1251 | 1.54 | 0.2143 |
| SA45 symptom severity | 1 | 1251 | 3.89 | 0.0489 | 1 | 1251 | 4.72 | 0.0299 |
| Psychosis | 1 | 1251 | 13.55 | 0.0002 | 1 | 1251 | 24.73 | <.0001 |
| Mania | 1 | 1251 | 0.65 | 0.4192 | 1 | 1251 | 1.37 | 0.2428 |
| Depression | 1 | 1251 | 7.60 | 0.0059 | 1 | 1251 | 4.81 | 0.0284 |
| DX:Sex | 3 | 1251 | 0.46 | 0.7134 | 3 | 1251 | 0.14 | 0.9376 |
| DX:Domain | 21 | 7762 | 3.60 | <.0001 | 24 | 8957 | 6.45 | <.0001 |
| Sex:Domain | 7 | 7762 | 4.58 | <.0001 | 8 | 8957 | 8.69 | <.0001 |
| DX:Sex:Domain | 21 | 7762 | 1.33 | 0.1428 | 24 | 8957 | 1.92 | 0.0043 |

**Supplementary Table 6. "Uni" is the uniqueness for each symptom from the 3-factor solution.** Factor correlations are presented below the loadings

| Symptom | **Uni** |
|---|---|
| Auditory Hallucinations | 0.29 |
| Delusion of Being Controlled | 0.36 |
| Visual Hallucinations | 0.42 |
| Bizarre Delusion | 0.33 |
| Tactile Hallucinations | 0.66 |
| Delusion of Reference | 0.36 |
| Persecutory Delusion | 0.26 |
| Grossly Disorganized Behavior | 0.31 |
| Other Delusions | 0.54 |
| Disorganized Speech | 0.4 |
| Religious Delusion | 0.5 |
| Suicidal thoughts lifetime | 0.38 |
| Suicide attempt | 0.68 |
| Depressed Mood | 0.24 |
| Anhedonia | 0.3 |
| Fatigue | 0.38 |
| Hypersomnia | 0.94 |
| Flight of Ideas | 0.37 |
| Decreased Need for Sleep | 0.33 |
| Grandiosity | 0.37 |
| Avolition | 0.62 |

**Correlations among factors**

| | **Psychosis** | **Depression** | **Mania** |
|---|---|---|---|
| **Psychosis** | 1 | -0.61 | 0.28 |
| **Depression** | -0.61 | 1.0 | -0.35 |
| **Mania** | 0.28 | -0.35 | 1 |

**Supplementary Table 7. Relationship between WAT, number of hospitalizations and ER visits, or SA45 global severity index, and Factor Scores or DX.** In models with Diagnosis, MDD is the reference.

| **A) factor scores** | **WAT** | | | | **number of hospitalizations and ER visits** | | | | **SA45 severity** | | | |
|---|---|---|---|---|---|---|---|---|---|---|---|---|
| | **Estimate** | **SE** | **t** | **Pr(>\|t\|)** | **Estimate** | **SE** | **t** | **Pr(>\|t\|)** | **Estimate** | **SE** | **t** | **Pr(>\|t\|)** |
| (Intercept) | 30.429 | 0.2135 | 142.538 | <2e-16 | 7.6552 | 0.2748 | 27.861 | <2e-16 | 43.234 | 1.017 | 42.531 | <2e-16 |
| Psychosis | -1.3073 | 0.2696 | -4.849 | 1.35E-06 | 2.4366 | 0.3672 | 6.636 | 4.69E-11 | 5.705 | 1.349 | 4.23 | 2.50E-05 |
| Mania | 0.5754 | 0.229 | 2.512 | 0.0121 | 2.4832 | 0.2967 | 8.369 | <2e-16 | -6.246 | 1.096 | -5.701 | 1.48E-08 |
| Depression | 0.5742 | 0.2772 | 2.071 | 0.0385 | 0.0392 | 0.366 | 0.107 | 0.915 | 11.572 | 1.356 | 8.536 | <2e-16 |
| **B) diagnosis + factor scores** | | | | | | | | | | | | |
| | **Estimate** | **SE** | **t** | **Pr(>\|t\|)** | **Estimate** | **SE** | **t** | **Pr(>\|t\|)** | **Estimate** | **SE** | **t** | **Pr(>\|t\|)** |
| (Intercept) | 31.0897 | 0.4213 | 73.79 | <2e-16 | 5.3199 | 0.5606 | 9.489 | <2e-16 | 49.697 | 2.058 | 24.153 | <2e-16 |
| BP2 | -0.5601 | 0.829 | -0.676 | 0.49937 | -0.4471 | 1.0746 | -0.416 | 0.677388 | 6.722 | 3.999 | 1.681 | 0.093 |
| BP1 | -1.8966 | 0.831 | -2.282 | 0.02259 | 4.8816 | 1.1154 | 4.377 | 1.30E-05 | -17.231 | 4.107 | -4.196 | 2.91E-05 |
| SCZ | -0.1169 | 1.1137 | -0.105 | 0.91643 | 10.2865 | 1.4755 | 6.972 | 4.93E-12 | -22.487 | 5.446 | -4.129 | 3.88E-05 |
| Psychosis | -1.3238 | 0.3186 | -4.155 | 3.41E-05 | 0.8605 | 0.423 | 2.034 | 0.042116 | 9.249 | 1.55 | 5.966 | 3.15E-09 |
| Mania | 1.2337 | 0.3845 | 3.209 | 0.00136 | 2.0154 | 0.5352 | 3.766 | 0.000173 | -3.596 | 1.964 | -1.831 | 0.0674 |
| Depression | 0.5769 | 0.2993 | 1.928 | 0.05406 | 0.9963 | 0.3881 | 2.567 | 0.01037 | 9.047 | 1.443 | 6.269 | 4.98E-10 |
| **C) diagnosis** | | | | | | | | | | | | |
| | **Estimate** | **SE** | **t** | **Pr(>\|t\|)** | **Estimate** | **SE** | **t** | **Pr(>\|t\|)** | **Estimate** | **SE** | **t** | **Pr(>\|t\|)** |
| (Intercept) | 31.0598 | 0.3111 | 99.823 | <2e-16 | 3.7121 | 0.3824 | 9.707 | <2e-16 | 52.478 | 1.446 | 36.279 | <2e-16 |
| BP2 | 0.7932 | 0.6913 | 1.147 | 0.2514 | 2.2002 | 0.843 | 2.61 | 0.00916 | 2.994 | 3.234 | 0.926 | 0.355 |
| BP1 | -1.1567 | 0.4975 | -2.325 | 0.0202 | 8.4996 | 0.6265 | 13.567 | <2e-16 | -24.303 | 2.362 | -10.29 | <2e-16 |
| SCZ | -3.934 | 0.7653 | -5.14 | 3.07E-07 | 10.9701 | 1.0239 | 10.714 | <2e-16 | -20.458 | 3.87 | -5.287 | 1.47E-07 |
